# JNK activation in TA and EDL muscle is load-dependent in rats receiving identical excitation patterns

**DOI:** 10.1101/2021.02.04.429746

**Authors:** Einar Eftestøl, Martino V. Franchi, Stephanie Kasper, Martin Flück

**Affiliations:** Department of Biosciences, University of Oslo, Norway; Laboratory for muscle plasticity, Department of Orthopaedics, University of Zürich, Switzerland; Department of Biomedical Sciences, University of Padova, Italy

**Keywords:** Exercise, hypertrophy, load, mechanotransduction, phosphorylation, skeletal muscle

## Abstract

**Aim:** As the excitation-contraction coupling is inseparable during voluntary exercise, the relative contribution of the mechanical and neural input is poorly understood. Herein, we use a rat in-vivo strength training setup with an electrically induced standardized excitation pattern previously shown to lead to a load-dependent increase in myonuclear number and hypertrophy, to study acute effects of load per se on molecular signalling.

**Methods:** Anaesthetized rats were subjected to unliteral identical electrically-paced contractions of the TA and EDL muscles under a high or low load for a duration of 2, 10 or 28-minutes. Muscle soluble proteins were extracted, and abundance and specific phosphorylations of FAK, mTOR, p70S6K and JNK were measured. Effects of exercise, load, muscle and exercise duration were assessed.

**Results:** Specific phosphorylation of S2448-mTOR, T421/S424-p70S6K and T183/Y185-JNK was increased after 28-minutes of exercise under the high- and low-load protocol. Elevated phosphorylation of mTOR and JNK was detectable already after 2 and 10 minutes of exercise, respectively, but greatest after 28-minutes of exercise. T183/Y185-JNK and S2448-mTOR demonstrated a load-dependent increase in phosphorylation in the exercised muscles that for mTOR depended on muscle type. The abundance of all four kinases was higher in TA compared to EDL muscle. FAK and JNK abundance was reduced after 28 minutes of exercise in both the exercised and control muscle.

**Conclusion:** The current study shows that JNK and mTOR activation is load-driven, and together with muscle-type specific mTOR and p70S6K effects it may drive muscle-type specific exercise and load-responses.

## INTRODUCTION

The biomechanical stress produced in a contracting muscle is believed to activate biochemical^1^ and biophysical^2^ signalling pathways, termed mechanotransduction,^3,4^ leading to increased biosynthetic efficiency and capacity that in the long-run result in functional muscle growth.^5^ A de novo hypertrophic response is also preceded^6^ and determined^7^ by myonuclear accretion,^8,9^ further contributing to increased synthetic capacity.

Although there is scarce experimental evidence for the underlying mechanisms, biomechanical stress is widely accepted as a primary trigger for muscle hypertrophy,^10–12^ and the effect of e.g. voluntary strength training or synergist-ablation induced overload on immediate-early signalling cues involved in protein synthesis and/or the long-term hypertrophic response is, by and large, implicated in mechanotransduction.^13^ However, under the above-mentioned conditions, as well as under most other experimental conditions, the neural input changes concomitantly, as an increased firing frequency and motor unit recruitment orchestrates the magnitude of force output and the following biomechanical stress.^14–19^ Thus, in experimental conditions that do not separately control neural input and mechanical stimuli it is difficult to properly address the importance of mechanotransduction in hypertrophy; it follows that the intrinsic nature of the molecular pathways that are implicated in the mechano-induced activation of muscle protein synthesis are still poorly characterized and understood^20–22^.

A well characterized route of activation of skeletal muscle protein synthesis (and subsequent hypertrophy) in response to resistance exercise is regulated by changes in the translational efficiency and capacity via signalling through coupled kinases, and is further increased through insertion of satellite cells.^23–26^ This increased translational capacity following resistance exercise in humans is correlated with increased muscle mass.^27–29^ The non-receptor tyrosine kinase FAK (focal adhesion kinase), and the downstream serine/threonine phosphotransferases mTOR (mammalian target of rapamycin) and p70S6K (70 kDa ribosomal protein S6 kinase) is implicated in the load-induced hypertrophic response in cell culture^30^ and after reloading of unloaded rats.^31^ Further, the post-translational modification and subsequent activation of FAK and the functionally-associated mixed-lineage kinase JNK (c-jun N-terminal kinase), as well as mTOR and p70S6K, correlate with the mechanical load impact during acute electrically-paced exercise.^32–34^ FAK and JNK have been regarded as part of the load-regulated myogenic adaptations in skeletal muscle.^35,36^ Especially, we have found that quantitative increases in the abundance and Y397 phosphorylation of FAK act upstream of muscle loading-related phosphorylation of T183/Y185-JNK and downstream of S63-c-jun and T421/S424-p70S6K.^37,38^ Interestingly, these mechanical pathways are also suggested to act synergistically with changes in amino acid uptake and growth factor signalling to sustain mTOR and FAK signalling.^21,39,40^ Thus, signalling pathways activated by e.g. mechano-, nutrient- and growth factor-stimuli probably modulate each other rather than working as independent pathways.

A few experimental studies have to a certain extent specifically addressed acute^32–35,41,42^ or long-term^34,43^ effects of mechanical stimuli per se, either by passively stretching innervated,^33^ (functionally) denervated^44–46^ or isolated^35^ muscles, or by standardizing the neural input by electrically-paced contractions either in vivo^32–34,41–43,47^ or in isolated muscles.^48,49^ However, a more comprehensive mapping and understanding of the activation of early mechanotransduction pathways connected to long-term outcomes are still warranted. To start on this quest, we previously developed a rat strength-training model and found that after a 6-week training regime under identical neural activity, differences in mechanical conditions had a major effect on the number of myonuclei and fibre size but not on fibre type.^22^ Also, after one acute session a few molecular correlates were indicated in the mechano-response, e.g. myogenin and MRF4, implicated both in satellite cell activation and an increased transcriptional response. Yet, no significant effect on the previously proposed mechano-responsive kinases Akt and p70S6K was observed.^22^

Thus, herein we investigate in more detail the acute responses to a single bout of the previously characterized training-model^22^ related to the proposed mechanosensitive and kinases FAK, mTOR, p70S6K and JNK. We find that JNK signalling is highly sensitive to mechanical stimuli, and propose a potential role for all 4 kinases in the following mechano-dependent hypertrophic response.

## RESULTS

A single exercise bout/session of electrically-paced contractions was evaluated for effects on specific phosphorylation (phosphorylated protein / total protein abundance) and total protein abundance for the kinases FAK, mTOR, p70S6K and JNK. 50 rats were unilaterally exercised for a total time of 2 (N=16), 10 (N=18) or 28 (N=16) minutes by stimulating the dorsiflexor muscle group (TA and EDL) non-invasively through skin-electrodes. Groups were equally allocated between a high-load or a low-load contraction, contralateral muscles functioning as non-stimulated controls. The low-load group obtained a peak tension of 50-60% of the high-load group.

### Effects of a single high- or low-load exercise bout on the phosphorylation of FAK, mTOR, p70S6K, and JNK in the EDL and TA muscle

A single 28-minute-long exercise stimulus led to an increased specific phosphorylation of S2448-mTOR (p<0.001), T421/S424-p70S6K (p<0.001), and T183/Y185-JNK (p<0.001), but not Y397-FAK (p=0.8), in both the TA and EDL muscle for both the low- and high-load exercised group compared to contralateral non-trained controls (main effect). A main interaction effect of exercise x load was found for the specific phosphorylation of T183/Y185-JNK (p=0.002), S2448-mTOR (p=0.025) and T421/S424-p70S6K (p=0.004). For JNK this effect was localized to an increased phosphorylation in the exercised leg after the high-load compared to the low-load exercise for both EDL (p=0.0085) and TA (p=0.0097) (Figure 1D). No such load-effect was observed at the post-hoc level for mTOR (Figure 1B) or p70S6K (Figure 1C).

**FIGURE 1.**
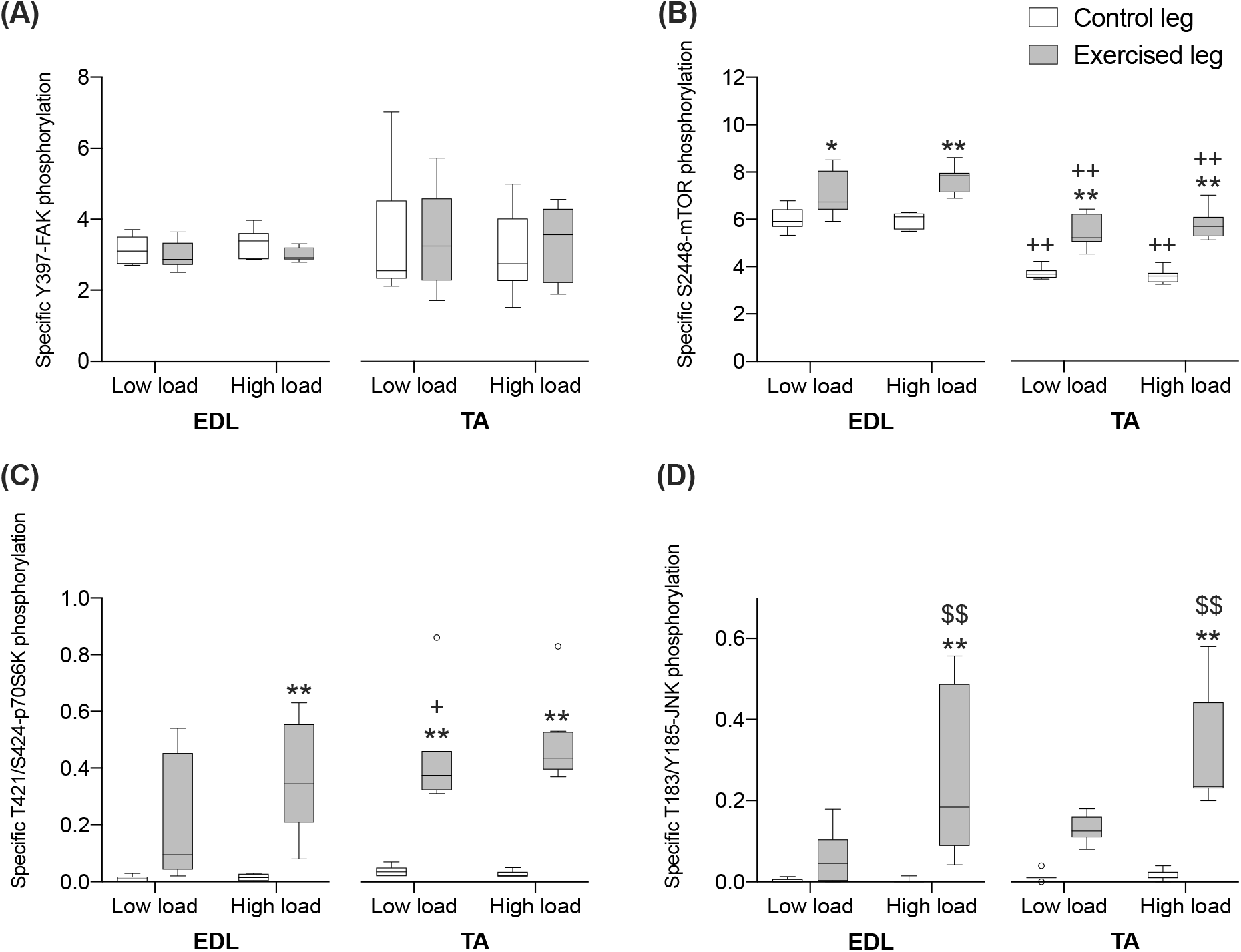
Exercise-induced JNK phosphorylation is load-dependent, while mTOR and p70S6K phosphorylation is muscle-type dependent. Specific phosphorylation of Y397-FAK (A), pS2448-mTOR (B), pT421/S424-p70S6K (C), and pT183/Y185-JNK (D) in exercised and contralateral control TA and EDL muscles after a 28-minutes exercise stimulus with a high or a low load. Significant differences denoted with *, ** p < 0.05 and 0.01 vs control leg; +, ++ p < 0.05 and 0.01 vs EDL within same exercise group. $$ p < 0.01 vs low load exercised group within the same muscle. Group data are presented as Box and whiskers plots (Tukey) (n=8 per group).

### Muscle-type differences in specific kinase phosphorylation

A main effect of the muscle type was observed for the specific phosphorylation of S2448-mTOR (p<0.001) and T421/S424-p70S6K (p=0.015), mTOR being higher in EDL when compared to corresponding group in TA (p<0.0001) (Figure 1B), and p70S6K being lower in EDL than TA (p=0.047) (Figure 1C).

### Muscle-type-, exercise- and load-effects on kinase abundance

The protein abundance of all assessed protein kinases demonstrated a main effect of the muscle type. FAK (p=0.002), JNK (p<0.001), and p70S6K (p<0.001) protein abundance was ~2-times higher and mTOR (p<0.001) abundance was ~10-times higher in TA than EDL muscles (Figure 2).

**FIGURE 2.**
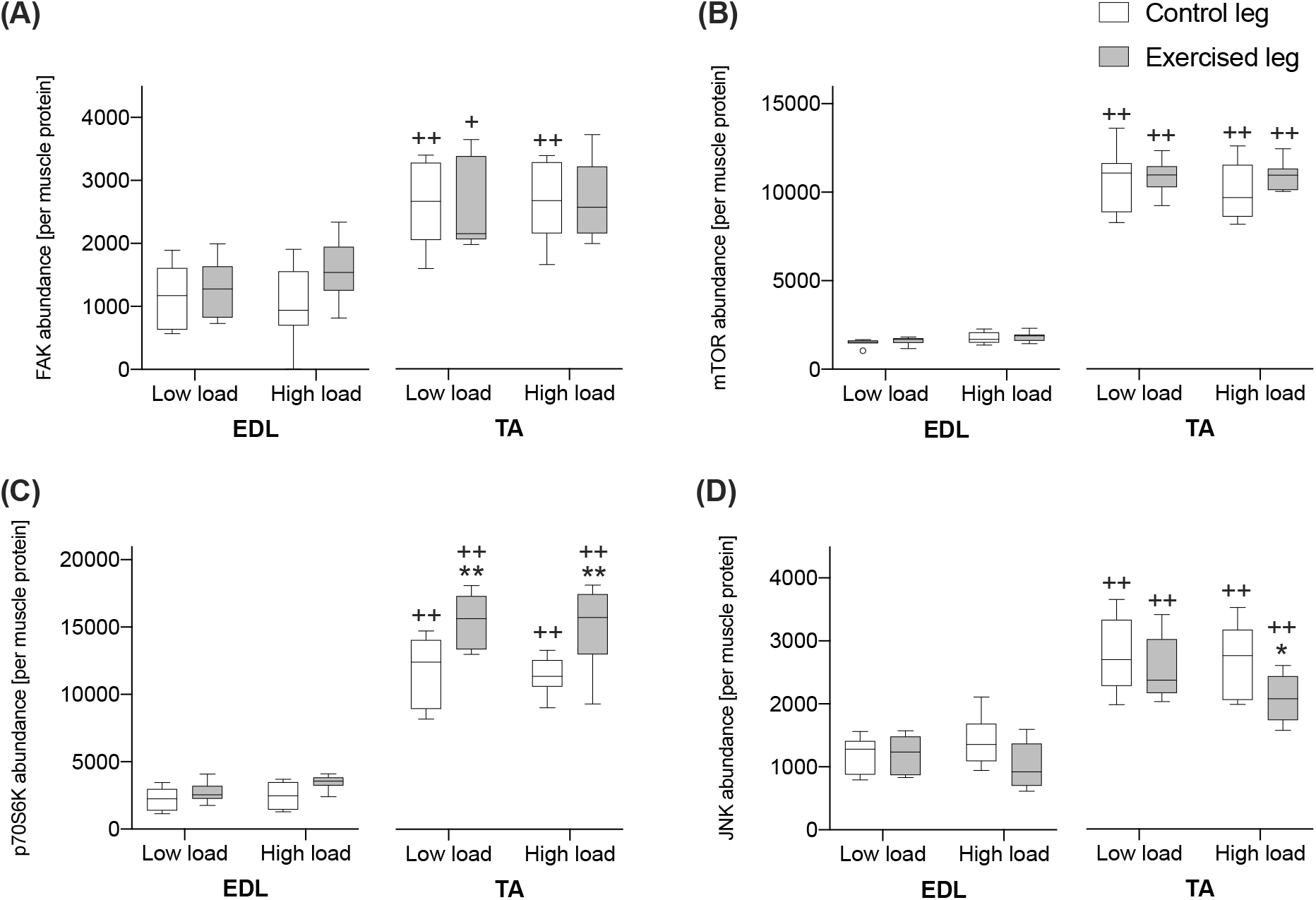
Kinase abundance is dependent on muscle type, and p70S6K abundance is increased by exercise in TA, while JNK abundance is decreased by exercise in a load-dependent manner in TA. Kinase abundance of FAK (A), mTOR (B), p70S6K (C), and JNK (D) in exercised and contralateral control TA and EDL muscles after a 28-minutes exercise stimulus with a high or a low load. Significant differences denoted with *, ** p < 0.05 and 0.01 vs control leg; ++ p < 0.01 vs EDL within same exercise group. Group data are presented as Box and whiskers plots (Tukey) (n=8 per group).

The abundance of P70S6K demonstrated a main effect of exercise (p<0.001), and an exercise x muscle interaction effect (p=0.006). These effects were localized to an increased abundance of p70S6K in exercised compared to control muscle for both low load (p=0.007) and high load (p=0.007) in TA, but not in EDL (Figure 2C).

The abundance of JNK demonstrated a main effect of exercise (p=0.032), and there was an interaction effect between exercise x load (p=0.015), and load x muscle type (p=0.028) for JNK abundance. This is explained by a reduced JNK abundance in 28-minutes exercised muscle after high-load exercise in TA (p=0.041), but not after low-load exercise (Figure 2D).

### Effects of exercise duration on the specific kinase phosphorylation in TA muscle

We further assessed whether the increase in the specific phosphorylation of mTOR, JNK (and FAK) in TA muscle after 28 minutes of exercise would be evident also at earlier time-points, i.e., after 2 or 10 minutes of exercise, and whether it would be load-dependent. The specific phosphorylation of S2448-mTOR and T183/Y185-JNK (both p<0.0001), but not Y397-FAK (p=0.57), demonstrated a main effect of exercise and also an interaction effect of exercise x exercise duration (S2448-mTOR (p=0.0003); T183/Y185-JNK (p<0.0001)). The specific phosphorylation of T183/Y185-JNK, alone, demonstrated an interaction effect of exercise x exercise duration x load (p<0.0006).

At the post hoc level, the specific phosphorylation of S2448-mTOR in TA muscle was increased in the exercised compared to the control leg after all durations of exercise, i.e., after 2, 10 and 28 minutes of exercise, in both the low-load and high-load group (for all comparisons p<0.001) except for after 2 minutes in the high-load group (Figure 3B). Specific phosphorylation of T183/Y185-JNK in TA muscle was increased in the exercised compared to the control leg after 10 (p=0.028) and 28 minutes (p<0.0001) of exercise in the high-load group, but first after 28 minutes in the low-load group (p<0.0002) (Figure 3C).

**FIGURE 3.**
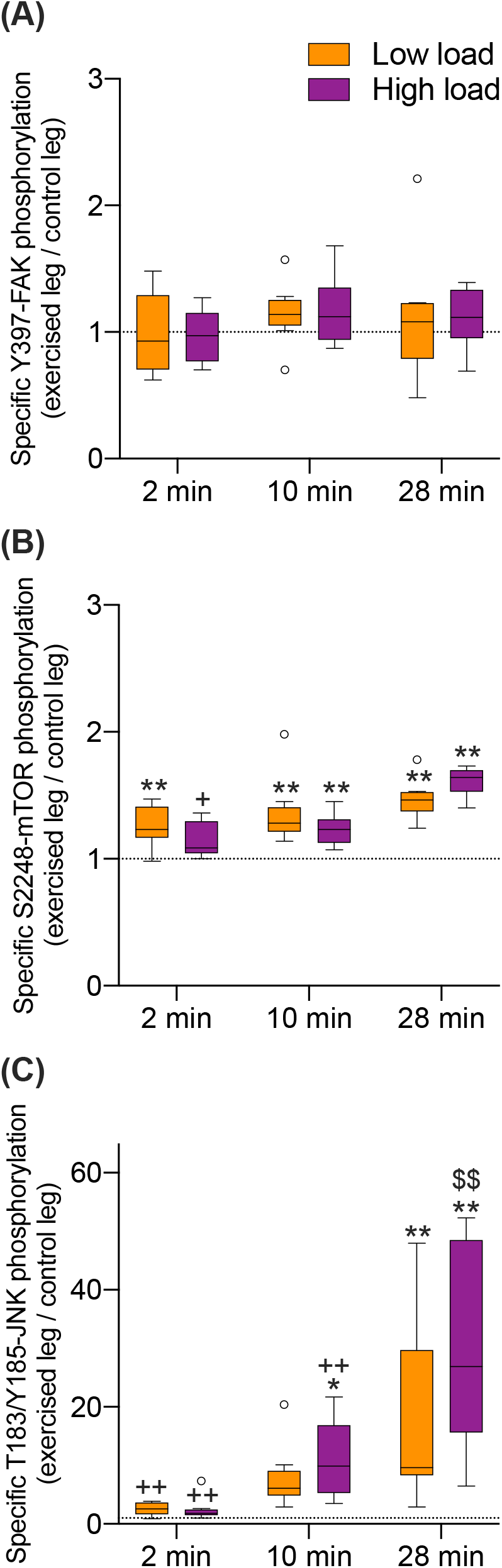
mTOR and JNK phosphorylation is increased with exercise duration, and for JNK also in a load-dependent manner. Specific phosphorylation of Y397-FAK (A), pS2448-mTOR (B) and pT183/Y185-JNK (C) in exercised relative to their respective contralateral control TA muscles after 2,10 and 28 minutes of exercise with a high or a low load. Significant differences denoted with *, ** p < 0.05 and 0.01 vs control leg; +, ++ p < 0.05 and 0.01 vs 28 minutes exercise within loading groups. $$ p < 0.01 vs low load exercised group after 28 minutes of exercise. Group data are presented as Box and whiskers plots (Tukey) (n=8 per group). Dotted line at y=1 representing no difference between exercised leg and contralateral control leg.

The specific phosphorylation of S2448-mTOR was increased after 28 minutes relative to 2 minutes of high-load (p=0.016), but not low-load exercise (Figure 3B). For T183/Y185-JNK, specific phosphorylation was increased after 28 minutes relative to 2 and 10 minutes of high-load (both p<0.0001), but only relative to 2 minutes of low-load (p=0.0007) exercise (Figure 3C). The specific phosphorylation of T183/Y185-JNK was also increased in the high-load vs low-load group after 28 minutes of exercise (p<0.0001), but not after 2- or 10 minutes (Figure 3C).

### Effect of exercise duration on kinase abundance in TA muscle

We further assessed whether alterations in protein abundance contributed to the observed effects of exercise on the specific kinase phosphorylation in TA muscle. No effect of exercise nor load was observed for any of the kinases, but a main effect of exercise duration was identified for the abundance of FAK (p=0.0014) and JNK (p=0.0008). Compared to the 2-minute and 10-minute duration of exercise, the abundance of FAK (p<0.004) and JNK (p<0.002) in TA muscle was lower after 28-minutes of exercise (Figure 4).

**FIGURE 4.**
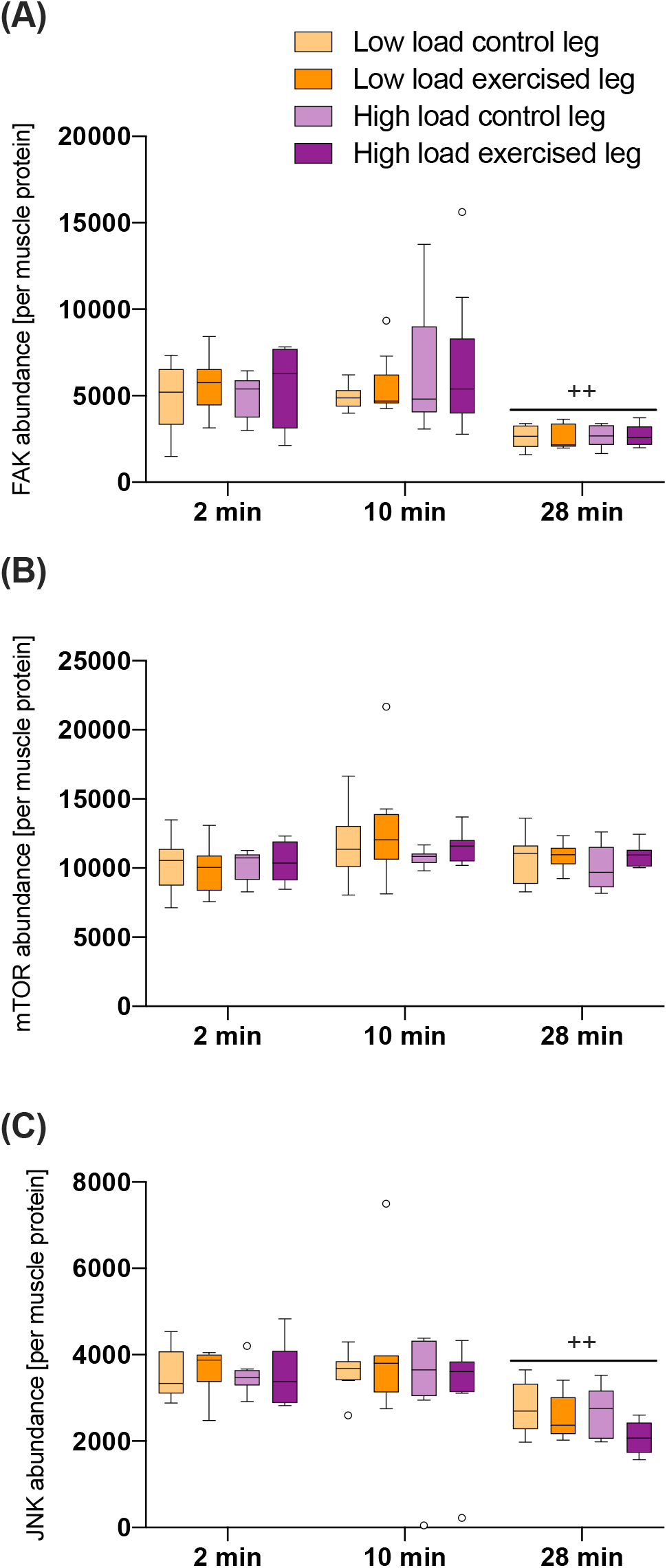
FAK and JNK abundance is decreased with exercise duration in both exercised and contralateral control leg, but not by exercise nor load per se. Kinase abundance of FAK (A), mTOR (B) and JNK (C) in exercised and contralateral control TA muscles after 2, 10 and 28 minutes of exercise with a high or a low load. Significant differences denoted with ++ p < 0.01 vs 2- and 10-minutes exercise irrespective of group. Group data are presented as Box and whiskers plots (Tukey) (n=8 per group).

## DISCUSSION

The relationship between mechanical signalling following a single exercise stimulus and long-term adaptations under conditions where the neuronal input is standardized is poorly characterized. We here used an electrically-paced, in-vivo rat strength training setup which has previously been shown to induce myonuclear accretion and hypertrophy^22^ in order to investigate early effects of mechanical load on the activation of FAK, mTOR, p70S6K and JNK, being implicated in the pre-translational regulation of protein synthesis, and previously suggested to be involved in mechanotransduction. We show here that JNK activation is highly load-dependent, and propose that observed effects of exercise, load, exercise duration and muscle-type on JNK, mTOR and p70S6K phosphorylation and abundance further our understanding of how the hypertrophic response is consolidated.

### JNK is activated by mechanical stimuli per se

We found JNK to be phosphorylated and thereby activated in a load-dependent manner in both TA and EDL. This is supported by an exercise-duration dependent effect already after 10 minutes of exercise only in the high-load group, increasing further after 28 minutes, and finally having increased JNK phosphorylation in the high-load group compared to the low-load group after the full 28-minute exercise session. An interaction effect of exercise and load was also evident for mTOR and p70S6K. To our knowledge, the present results, for the first time, link acute specific effects of load on JNK activation with previously observed long-term hypertrophic effects of the same exercise stimulus.^22^

In rat plantaris muscle, an increased JNK phosphorylation has been shown to correlate with mechanical stress after a single bout of in situ eccentric (~10% length change) compared to concentric and isometric electrically-paced contractions.^33^ JNK activation is further shown to be required in synergist ablation-induced muscle hypertrophy^35^ and this involves the counteraction of myostatin signals which inhibit the AKT-mTOR pathways and the upregulation of ubiquitin-mediated proteasomal degradation.^50^ These effects seem also to involve the activation of Jun expression which is elevated by muscle stretch^51^ in interstitial and resident myonuclei^52^ that may upregulate the promoter activity of muscle genes via Jun/Fos-heterodimers (e.g. AP-1^53,54^). Nevertheless, the observed exercise-induced specific phosphorylation of JNK is likely not sufficient to explain the muscle- and load-dependent degree of hypertrophy observed in a previous study by Eftestøl et al.^22^ after repeated impact of the same 28-minute protocol as herein over a period of 6 weeks. For instance, larger increases in the cross-sectional area of muscle fibres were reported with the high-load compared to the low-load protocol in the TA muscle (i.e. 33% vs 18%) compared to the EDL muscle (i.e. 16% vs 0%).^22^ The consistently higher abundance of all assessed protein kinases in TA compared to EDL combined with the muscle-type specific effects on mTOR and p70S6K phosphorylation observed herein reflects the previously observed higher degree of hypertrophy in TA compared to EDL muscle.^22^ This correspondence suggests that quantitative differences in the expression of the signalling molecules mTOR and p70S6K are part of the mechanism that sets the overall pace of the hypertrophic response. In addition, JNK activation could be related to other aspects of the observed load-dependent muscle responses such as the increased myonuclear number (i.e., satellite cell activation), as well as increased transcript expression of myogenin and myogenic regulatory factor 4.^22^

Interestingly we observed an increased activation of both mTOR and p70S6K with exercise, including an interaction effect with load, yet not significantly so by load at the post-hoc level. An important aspect of the present in-vivo setup is that due to the physical limits of the joint excursion-range, the peak tension in the low-load group could not be decreased to less than about 50-60% of the corresponding high-load group, as that would require an even faster concentric contraction, resulting in a high-load isometric contraction towards the end of the stimulation. Notable, after 28 minutes of exercise, there was a consistently higher degree of phosphorylation (~10%) in the high-load vs low-load group for mTOR and p70S6K in both TA and EDL, which is also in line with previous observations for Akt and p70S6K phosphorylation.^22^ It is thus intriguing to speculate that with a larger difference between the loading groups, all these kinases, that are widely known to be important in regulating protein synthesis (see introduction), would reveal a load-dependent activation. In fact, Rindom et al.^48^ has recently shown that, in response to acute exercise by electrically stimulating isolated rat EDL muscles, mTOR and p70S6K phosphorylation was load-dependent, but Jun and Fos mRNA transcription was dependent on both excitation and tension development.^48^ Importantly, such in vitro setups are not prone to the same limitations as in-vivo setups, as they are e.g. not limited by the joint excursion-range. Thus, through the use of chemical inhibitors of the muscle contraction together with passive stretch, Rindom et al.^48^ were able to compare a high-load active contraction to both a high-load passively stretched group with comparable tension development either with or without excitation, and also to an excited “low-load” group with ~100% reduction in tension development.

In summary, in light of the recent in-vitro work by Rindom et al.^48^, the in-vivo results presented herein strengthen the notion that mechanical stimuli per se, uncoupled from the active contraction, is an important signalling cue for protein synthesis, yet specific signalling cues from the excitation are important to achieve a full activation of protein synthesis and following hypertrophy. Further, JNK is activated by mechanical stimuli per se, further indicating that this is important for eliciting the load-induced hypertrophic response previously observed with long-term strength training.^22^ Also, it is likely that both mTOR and p70S6K are part of a mechanotransduction network in conjunction with JNK, modulated both by mechanical stimuli and other training-related stimuli^21,55^ like the excitation-contraction coupling and metabolic cues, all contributing to elicit a full hypertrophic response.

### Effects of exercise duration and load on FAK and JNK abundance

The abundance of FAK and JNK, but not mTOR, was affected to a considerable degree by the duration of the exercise experiment per se, which was explained by a reduced abundance in the assessed soluble protein fraction after the 28-minutes exercise bout in both the control and exercised leg. Stretch increases the phosphorylation of FAK in rat skeletal muscle within 20 seconds,^56^ and translocation of activated FAK from sarcomeres to a nuclei-associated, Triton X-100 insoluble, fraction has been reported to occur in the heart within 60 minutes of pressure overload, corresponding with increases in protein synthesis and myofibrillar assembly.^57^

Our present data thus indicate a possible experimental model-related interference with FAK and JNK signalling through effects on their solubility and/or calcium-induced degradation^58^ during the 28-minutes of exercise. Both FAK and JNK are part of a rescue pathway that can be conditioned by local ischemia to protect skeletal muscle from reperfusion injury,^59,60^ suggesting that systemic effects in the exercising and control muscle may be due to the experimental procedure, including, but not limited to, training-induced ischemia. In this respect, an altered solubility may have influenced the observed increase in the specific phosphorylation of JNK 28-minutes, but not 2 and 10-minutes, after exercise. That the JNK abundance in the assessed soluble fraction was lower in the high-load than the low-load group after the 28-minutes exercise thus strengthens our conclusion that JNK is preferentially activated by high-load, given the notion that it is the activated form that translocates to the insoluble fraction.^61^ Probably this relates to the activation of JNK in rat skeletal muscle by stretch or other mechanical cues.^33^ However, we cannot rule out a possible exercise- and load-effect also on FAK that we were not able to detect, given the addressed limitations of the experimental setup, related to the rapid activation- and translocation dynamics of FAK,^62^ as it has previously been indicated as an early mechanotransducing kinase in skeletal muscle.^1,37^

In summary, our study shows that JNK is specifically activated in a load- and time-dependent manner during in vivo strength training. It also furthers our understanding of the muscle-, time- and load-specific acute responses to electrically paced-exercise for the investigated kinases FAK, mTOR, p70S6K and JNK, and how these effects together could be linked to an increased protein synthesis and hypertrophic response.

## MATERIALS AND METHODS

### Design

The dorsal flexors, extensor digitorum longus (EDL) and tibialis anterior (TA), of the right lower leg of anesthetized 10 weeks old male Sprague Dawley rats with a mean body weight of 413 grams were randomly subjected to electrically paced contractions under a high or low load during 2, 10 or 28 minutes. The left non-stimulated left leg served as control. 8-9 animals were used per protocol. Soluble protein from the excised muscles were subjected to electrochemiluminescence (ECL)-based immunoassays to quantify the abundance of FAK, mTOR, p70S6K and JNK, as well as the respective level of phospho-Y397-FAK, phospho-S2448-mTOR, phospho-T421/S424-p70S6K, and phospho-T183/Y185-JNK.

### Electrically paced exercise

Muscle contractions were electrically paced under identical excitation patterns to have the stimulated right lower leg perform controlled isometric (high load) or concentric (low load) tetanic contractions under inhalation anaesthesia (isoflurane) essentially as described before.^22^

The skin of the lower leg of anesthetized rats was shaved, and two platinum skin electrodes measuring 5 times 6 mm covered with electrode gel (Spectra 360; Parker Laboratories) were placed on the skin over the dorsiflexor muscle group (TA and EDL) of the right leg about 5 mm apart. The electrodes were connected to a pulse generator (Pulsar 6bp-a/s; FHC).

Exercise was conducted over 90, 360, or 900 short tetanic contractions during 2, 10 or 28 minutes, each contraction lasting about 120 ms. Stimulation was delivered in trains of 10 consecutive 0.5 ms symmetrical bipolar pulses delivered at an instantaneous frequency of 150 Hz at amplitude 30-45 V. Each pulse train was delivered every 600 ms repeated 30 times. For the 2 minutes protocol, each such series was repeated every 40 s three times. For the 10 minutes protocol, each series was repeated every 40 s six times, and each such session was delivered every 360 s two times. For the 28 minutes protocol, each session was repeated five times.

During training, the right leg was attached to a footplate connected to a DC motor (Dual lever-arm system 305C-LR connected to a 600A Digital Controller; Aurora Scientific, Aurora, ON, Canada) in length-control mode, allowing the load to be differentiated during contraction. The high-load group performed pure isometric contractions while the low-load group performed mainly high velocity (550°/s) concentric contractions with 50-60% of the peak force produced in the high-load group. The exercise was performed during the rats’ dark cycle, and different groups were always exercised alternately to minimize circadian differences in training response. The foot was positioned at an angle of 90° relative to the tibia during the contraction in the high-load group while the low-load group moved from 128.5 to 90°. EDL and TA from the stimulated (i.e. exercising) right leg and then the non-stimulated (i.e. control) left leg was harvested in sequential order immediately after the exercise and snap frozen in liquid nitrogen as described earlier.^22^

### Protein extraction

Homogenate was prepared from pooled 20-micrometer-thick cryosection in ice-cold RIPA buffer (~3 mm3 volume per 200 microliters of 2% Triton X-100, 1% NP-40, 300mM NaCl, 20mM Tris base, 2mM EDTA, 2mM EGTA) with the help of a rotor-stator mixer (Polytron PT1200, KINEMATICA. AG, Lucerne, Switzerland). The soluble fraction of proteins recovered from the supernatant and protein concentration determined as described.^37^

### ECL-based immunoassay

A volume of supernatant corresponding to 33 micrograms of soluble muscle protein was analysed per sample for the abundance of p70S6K and phospho-T421/S424-p70S6K, mTOR and phospho-S2448-JNK, JNK and phosphor-T183/Y185-JNK, FAK and phospho-Y397-FAK with a validated MESO scale system (K15114D-1, K15170D-1, K15111D-1, or customized U-PLEX development system; Meso Scale Discovery, USA) as described.^37^

One sample each from TA and EDL muscle of the 28-minutes of exercise, and TA muscle from the 2 and 10 minutes of exercise, was analysed in separate runs on the same ELISA plate. Aliquots from four different samples were run on both ELISA plates to scale the ECL signals for the kinase abundance and phosphorylation from the two runs to the average of the four signals on each of the respective plate. This procedure was carried out to control for inter-assay variability due to differences in sample handling on the day of the experiment and variability in the quantitative relationship between the commercial and custom-prepared MESO scale plates from different batches.

The values for the resulting ECL signal in counts were exported in Microsoft Excel for Windows 365 (Microsoft Corporation, Redmond, USA) and used to express the relative abundance of each kinase per muscle protein, and calculate the specific phosphorylation of each kinase from the ratio of the ECL signal for the phosphorylated kinase versus the ECL signal for the respective kinase abundance. The fold differences in the specific phosphorylation of a kinase with exercise was calculated from the ratio of the respective values for the specific phosphorylation between muscles from the exercised leg and its contralateral control.

### Statistics

Data analysis and graphing was carried out with Prism version 9.0.0 for macOS (Graphpad Software, LLC). Repeated measures ANOVAs were carried out on the ECL-based values for the kinase abundance and specific phosphorylation for the repeated factors exercise (control left leg, exercised right leg) and where applicable muscle (EDL, TA), and the not-repeated factor load (high load, low load) and duration of exercise (2, 10 or 28 minutes). Sphericity was assessed with Mauchly's test statistic (this assumption was never rejected).

Main effects (p-values) from the three-way ANOVA for the factors exercise, load and muscle type (corresponding to Figure 1-2) or exercise, load and exercise duration (corresponding to Figure 3-4) are presented in the results section. Post hoc effects were localized with a Tukey’s multiple comparisons test and are denoted in the figures. Significance was declared at p < 0.05. All outliers are included in statistics and shown according to Tukey in figures as open circles.

## Notes

The authors declare no conflicts of interest

### Competing Interest Statement

The authors have declared no competing interest.

